# The AusAB non-ribosomal peptide synthase in *Staphylococcus aureus* preferentially incorporates exogenous phenylalanine and tyrosine into the aureusimine natural products

**DOI:** 10.1101/2024.03.22.586303

**Authors:** Adriana Moldovan, Markus Krischke, Claudia Huber, Clara Hans, Martin J. Müller, Wolfgang Eisenreich, Thomas Rudel, Martin J. Fraunholz

**Affiliations:** Chair of Microbiology, Biocenter, University of Würzburg, Würzburg, Germany; Biocenter, Chair of Pharmaceutical Biology, University of Würzburg, Würzburg, Germany; Bavarian NMR Center - Structural Membrane Biochemistry, Department of Bioscience, School of Natural Sciences, Technical University of Munich, Germany

## Abstract

Non-ribosomal peptide synthases (NRPS) are modular multidomain enzymes, responsible for the biosynthesis of various secondary metabolites, in a mRNA-template independent manner. They are predominantly present in bacteria and fungi, where they synthesize a variety of products, including antibiotics, siderophores, toxins and signalling molecules. The human pathogen *Staphylococcus aureus* possesses one single NRPS, AusA, highly conserved in all sequenced *S. aureus* strains. AusA incorporates the aromatic amino acids (AAA) phenylalanine or tyrosine, as well as the branched-chain amino acids (BCAA) valine and leucine into three cyclic dipeptides collectively called aureusimines: phevalin, tyrvalin and leuvalin. By using targeted metabolomics, we found that AusA preferentially incorporates phenylalanine and tyrosine from an exogenous source into aureusimines, whereas the source of valine can be either endo- or exogenous.

Upon cultivation in a chemically defined medium (CDM) lacking phenylalanine, the amino acid was not incorporated into phevalin, despite *de novo* phenylalanine biosynthesis. Tyrosine production remained unaffected. Conversely, upon cultivation in medium lacking tyrosine, tyrvalin production was not detected, despite tyrosine *de novo* biosynthesis. Phevalin production, however, remained unaltered. By contrast, omission of valine in the culture medium not only resulted in de novo valine biosynthesis but also was accompanied by phevalin production.

To our knowledge, this is the first report of a selective incorporation of AAAs by a bacterial NRPS, which provides useful basis for linking bacterial cell metabolic status to the biosynthesis of secondary metabolites.

**IMPORTANCE:** Peptide and protein synthesis are fundamental processes in nature, which are largely mediated by the ribosomal machinery. An alternative pathway for peptide synthesis is non-ribosomal mRNA-template independent synthesis, performed by so-called non-ribosomal peptide synthases (NRPS). NRPSs are multi-enzyme complexes, which serve the simultaneous role of template and biosynthetic machinery. They are mostly found in bacteria and fungi and are responsible for the biosynthesis of many pharmacologically significant products, including antibiotics, anticancer compounds or immunosuppressants.

The human pathogen *S. aureus* possesses one such NRPS, AusA, which synthesizes three cyclic dipeptides termed “aureusimines” using the aromatic amino acids phenylalanine and tyrosine, and the branched-chain amino acid valine. Although the biological role of aureusimines remains unknown, AusA appears to play a role in the interaction of *S. aureus* with the host. In addition, owing to its minimal canonical NRPS structure and autonomous function (i.e. most NRPS pathways require the assembly of several NRPS proteins), AusA represents an excellent model system for studying such molecular assembly lines. Our observation is, to our knowledge, the first report of a NRPS incorporating phenylalanine and tyrosine only from exogenous sources (e.g. environment, culture medium), but not from *de novo* “self-made” pool. This opens up new avenues in understanding and modulating the function of NRPSs (e.g. for biotechnological purposes).

## INTRODUCTION

Non-ribosomal peptide synthases (NRPSs) are large, modular multi-enzyme complexes responsible for the production of complex peptide natural products with diverse properties such as antibiotics, anti-fungal reagents, siderophores, pigments or toxins [1]. They are widespread across all three domains of life, but gene clusters encoding NRPSs are more commonly reported in bacteria [2]. NRPSs employ a modular architecture wherein each module contains several domains responsible for introducing one amino acid into the building peptide. A canonical NRPS module contains three domains: an adenylation domain (A) responsible for the selection and activation of an amino acid, a thiolation domain (T), onto which the amino acid is covalently tethered as a thioester to the terminal thiol of 4’-phosphopantetheine prosthetic group, and a condensation domain (C) which catalyzes peptide bond formation. Terminal modules include domains dedicated to the release of the nascent peptide, either by the activity of a thioesterase domain (TE), via cyclization, or more rarely, by a reductase domain (R) [1, 3, 4]. Non-ribosomal peptide synthesis begins with an essential post-translational modification of the NRPS itself by a phosphopantetheine transferase (PPTase) which adds a 4’-phosphopanthetheine (Ppant) moiety to a conserved serine residue within the apo-T domains to yield functional holo-T domains [1, 5].

The genome of the prominent opportunistic pathogen *Staphylococcus aureus* encodes a single NRPS biosynthetic cluster, *aus*AB (SAUSA300_RS00950 and RS00955). AusA is a 277 kDa two-module six-domain NRPS, with the following domain architecture: A_1_-T_1_-C-A_2_-T_2_-R [4]. apo-AusA is post-translationally modified by the AusB PPTase (Fig. 1A).

**Fig 1.**
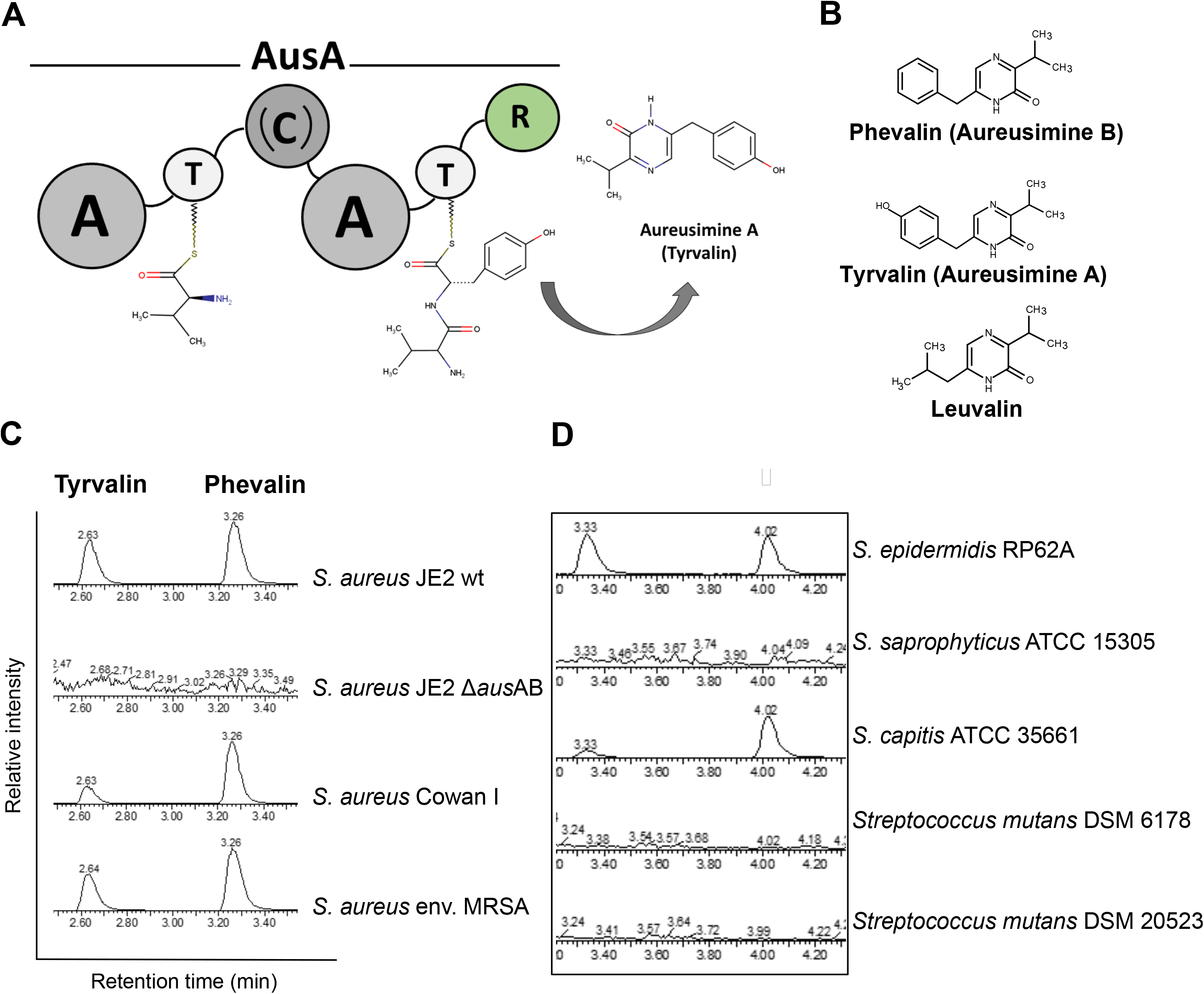
Aureusimines are synthesized by different *S. aureus* strains, and by skin-associated staphylococci. (**A**) Schematic representation of the AusA non-ribosomal peptide synthase (NRPS) (genomic locus ID: SAUSA300_RS0095). AusA is a ∼273 kDA soluble protein with the bimodular architecture: A_1_-T_1_-C-A_2_-T_2_-R (A=adenylation domain; T=thiolation domain; C=condensation domain; R=terminal reductase domain). (**B**) Aureusimines (phevalin, tyrvalin and leuvalin) are cyclic dipeptides with monoketopiperazine structure. (**C**) Phevalin and tyrvalin are synthesized by both cytotoxic (*S. aureus* JE2) and non-cytotoxic *S. aureus* (Cowan I), as well as by environmental *S. aureus*. (**D**) Phevalin and tyrvalin are detected in *S. epidermidis* and *S. capitis*, two skin-associated coagulase-negative staphylococcal species, but not in *Streptococcus mutans*. Phevalin and tyrvalin were detected by UPLC-MS in stationary phase culture supernatants of bacteria grown in TSB Medium for 24h. Commercially available phevalin and tyrvalin serve as standard. Leuvalin is to date not commercially available, therefore was not analyzed in the present study. Shown UPLC spectra are representative of two independent biological replicates (n=2).

AusA is responsible for the biosynthesis of three cyclic dipeptides, formed through the condensation of the aromatic amino acids (AAAs) L-phenylalanine or L-tyrosine and, to a lesser extent, L-leucine, with the branched-chain amino acid (BCAA) L-valine, to form three cyclic dipeptides with monoketopiperazine structure: phevalin, tyrvalin and leuvalin, collectively termed aureusimines [6, 7] (Fig. 1B).

The high degree of conservation of the *aus*AB gene cluster across all sequenced *S. aureus* strains, suggests an important biological function for the NRPS. Two studies propose a role for the AusAB for intracellular *S. aureus* virulence [8, 9]. Furthermore, the dipeptide aldehyde form of phevalin (i.e. the non-oxidized dihydropyrazinone, released by the C-terminal reductase domain of the NRPS) has been ascribed protease-inhibitory functions, in anaerobic gut microbiota species [10]. However, the exact function of any naturally occurring monoketopiperazine remains elusive.

We sought to gain insight into the biosynthesis of aureusimines under amino acid-depleted conditions, in order to mimic host niches where such conditions likely arise (e.g. human nasal secretions completely lacking tyrosine [11]).

## OBSERVATION

The *aus*AB locus is well conserved in all sequenced *S. aureus* strains. We therefore assessed by UPLC-MS several *S. aureus* strains for phevalin and tyrvalin production during growth in the complex medium TSB. Since *aus*AB genes were shown to be upregulated during stationary phase [8], we measured aureusimine production in the stationary phase of bacterial growth. The methicillin-resistant USA300 JE2 *S. aureus* strain, the non-cytotoxic, *agr-*negative strain Cowan I (ATCC 12598), as well as an environmental *S. aureus* strain were aureusimine producers (Fig. 1C, Supp. Fig. 1). AusA homologues of different identity levels were identified in the human-associated *S. epidermidis, S. capitis, S. schweitzeri, S. warneri* and *S. lugdunensis* [6]. While the degree of homology of the AusA protein was ∼53-54%, aureusimine production was detected in *S. epidermidis* and *S. capitis* (Fig. 1D; Supp. Fig. 2), as well as in *S. lugdunensis* (personal communication with Dr. Bernhard Krismer, Eberhard Karls Universität Tübingen). Neither *S. saprophyticus* ATCC 15305, nor two different *Streptococcus mutans* strains, a member of the human oral bacterial flora previously reported to synthesize phevalin [12] produced aureusimines under the tested conditions. Since *S. epidermidis, S. capitis* and *S. lugdunensis* are human-associated staphylococci [13-15], it is tempting to speculate that aureusimine production is associated with human commensal bacteria. This is supported by a study, which found phevalin, as well as several other cyclic dipeptides to be produced by NRPSs in human gut bacteria [10].

**Fig 2.**
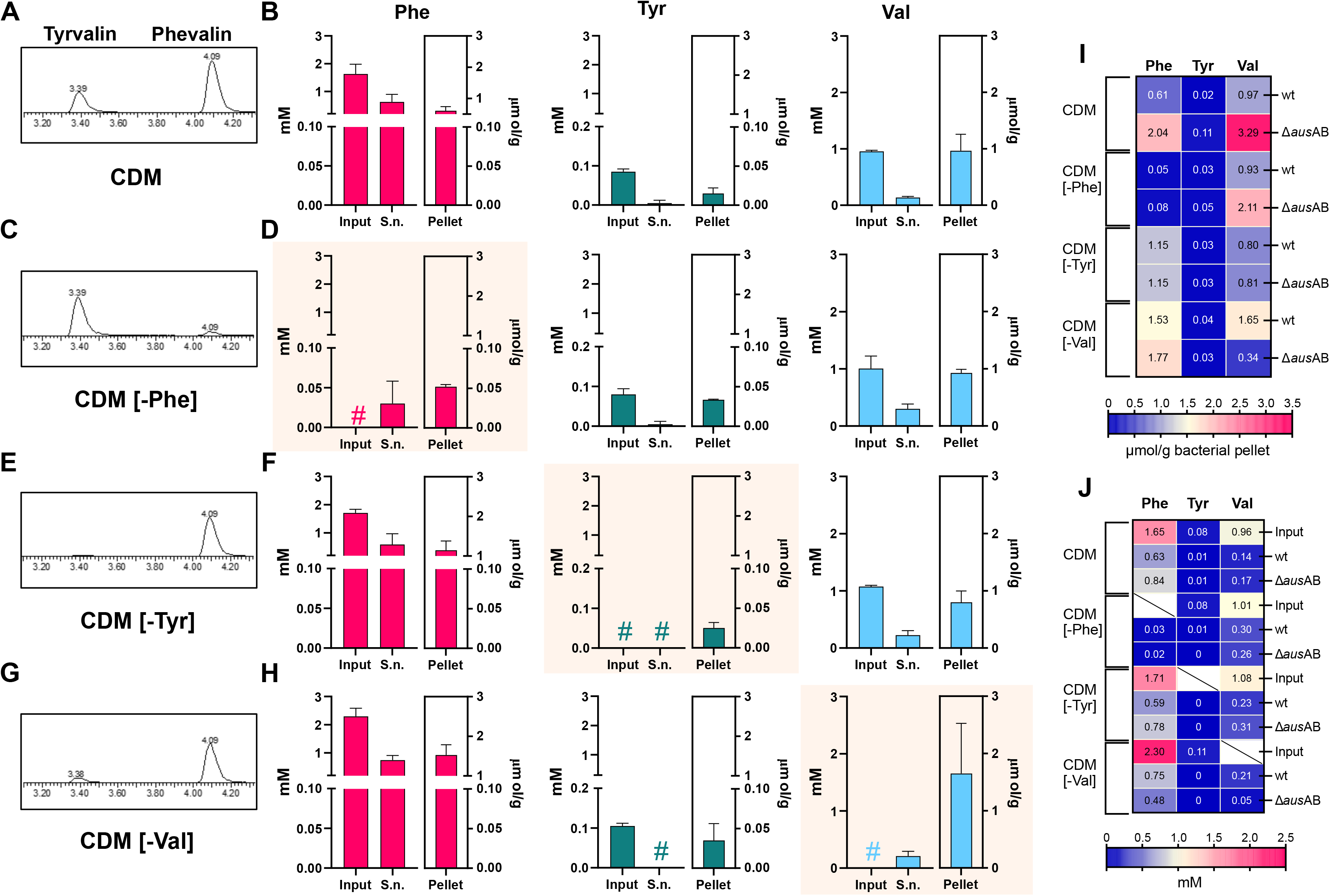
Aureusimine biosynthesis is dictated by exogenous aromatic amino acid availability in *S. aureus*. (**A**) Phevalin and tyrvalin detection by UPLC-MS in *S. aureus* JE2 wt supernatants grown in complete chemically defined medium (CDM) for 24h. (**B**) Detection of phenylalanine (Phe), tyrosine (Tyr) and valine (Val) in *S. aureus* JE2 cultures grown in CDM for 24h, either secreted (S.n.) or bacterial pellet-associated. (**C**) Tyrvalin detection by UPLC-MS in *S. aureus* JE2 wt supernatants grown in CDM without Phe (CDM [-Phe]) for 24h. (**D**) Detection of Phe, Tyr and Val in *S. aureus* JE2 cultures grown in CDM [-Phe] for 24h, either secreted (S.n.) or bacterial pellet-associated. (**E**) Phevalin detection by UPLC-MS in *S. aureus* JE2 wt supernatants grown in CDM without tyrosine (CDM [-Tyr]) for 24h. (**F**) Detection of Phe, Tyr and Val in *S. aureus* cultures grown in CDM [-Tyr] for 24h, either secreted (S.n.) or bacterial pellet-associated. (**G**) Phevalin detection by UPLC-MS in *S. aureus* JE2 wt supernatants grown in CDM without valine (CDM [-Val]) for 48h. (**H**) Detection of Phe, Tyr and Val in *S. aureus* cultures grown in CDM [-Val] for 48h, either secreted (S.n.) or bacterial pellet-associated. (**I**) Quantification of bacterial pellet-associated Phe, Tyr and Val for *S. aureus* JE2 wt and *S. aureus* Δ*aus*AB cultivated for 24h in complete CDM, CDM [-Phe] or CDM [-Tyr] and 48h in CDM [-Val] (data is expressed as μmol amino acid/g wet pellet) (**J**) Quantification of secreted Phe, Tyr and Val for *S. aureus* JE2 wt and *S. aureus* Δ*aus*AB cultivated for in complete CDM, CDM [-Phe] or CDM [-Tyr] (for 24h) or CDM [-Val] (for 48h h) (data is expressed in mM). Shown UPLC spectra (A, C, E and G) are representative of independent biological replicates (n=2). Commercially available phevalin and tyrvalin serve as standard. (B, D, F, H) Data are shown as mean +SD from independent biological replicates (n=2). Norvaline was used as internal standard for quantification. No statistical analysis was performed. (I, J) Heat-maps depict quantification data for Phe, Tyr and Val, using the same dataset as (B, D, F, H) for JE2 wt. Data are shown as mean from independent biological replicates (n=2). “#” denotes absence of the respective amino acid in the respective sample. Quantification of the amino acid omitted in the respective CDM formulation is highlighted.

Aureusimines are enigmatic natural products and, while AusAB appears to play a role in the virulence of intracellular *S. aureus* [8, 9], *the exact targets of these molecules are poorly understood. Naturally occurring phevalin was first isolated from Streptomyces* sp., and exhibited calpain inhibitory activity [16]. Synthetic phevalin was shown to inhibit human neutrophil activation and enhance phagosomal escape of *S. aureus* in epithelial cells [8], thus suggesting potential host targets for this molecule. We sought to gain insight into the biosynthesis of aureusimines under amino acid-depleted conditions, in order to mimic host niches where such conditions likely arise (e.g. human nasal secretions completely lacking tyrosine [11]). We therefore grew *S. aureus* JE2 in either a complete chemically defined medium (CDM) or CDM where phenylalanine, tyrosine or valine were excluded and analyzed the production of aureusimines under these conditions.

Upon cultivation in complete CDM, containing 1 mM L-phenylalanine and L-valine and 0.1 mM L-tyrosine, tyrvalin and phevalin were detected in stationary phase cultures (Figure 2A; Supp. Fig. 3). Since leuvalin is not commercially available and it represents a minor byproduct, we excluded it in this study. All three amino acids were present, both in bacterial supernatants and associated with the bacterial pellet (Figure 2B).

Upon exclusion of L-phenylalanine from the medium (CDM [-Phe]), expected amounts of tyrvalin, but only trace amounts of phevalin were detected (Figure 2C; Supp. Fig. 3). Phenylalanine was, however, present in both bacterial supernatants and pellets (Figure 2D), indicating *de novo* biosynthesis of the amino acid.

Conversely, upon exclusion of tyrosine from the CDM formulation (CDM [-Tyr]), phevalin, but not tyrvalin was detected in bacterial supernatants (Figure 2E; Supp. Fig. 3), despite *de novo* tyrosine biosynthesis by *S. aureus* being evidenced in the pellet-associated fraction, but not in the supernatant (Figure 2F).

Cultivation under valine depletion (CDM [-Val]) resulted in production of both aureusimines (Figure 2G; Supp. Fig. 3), albeit with notable reduction in tyrvalin biosynthesis. We confirmed *de novo* valine biosynthesis, as the amino acid was present in both, supernatant and the bacterial pellet (Figure 2H). Of note, bacterial growth pattern under either phenylalanine or tyrosine deprivation showed mild delay, compared to growth in complete CDM (Supp. Fig 4A and B). Consistent with previous reports [17], growth under valine deprived conditions was dramatically impacted, with *S. aureus* undergoing a 24h lag-phase before entering logarithmic growth (Supp. Fig 4C).

Quantification of the three amino acids in supernatants and cell fractions of both strains JE2 wt and JE2 Δ*aus*AB revealed only minor differences in the bacterial pellet-associated fraction, apart from valine which was detected in higher amounts in case of JE2 Δ*aus*AB in complete, as well as CDM [-Phe] media. In CDM [-Val], however, strikingly less valine was measured in JE2 Δ*aus*AB (Fig 2I). Similarly, measurements of bacterial supernatants revealed no prominent differences between JE2 wt and JE2 Δ*aus*AB, except for CDM [-Val], where lower amounts of valine were detected in case of JE2 Δ*aus*AB (Fig 2J).

Within the AusA NRPS, the A_1_ domain activates and loads T_1_ with L-Val, to yield L-Val-S∼T_1_; the A_2_ domain then activates and loads T_2_ with L-tyrosine to yield L-Tyr-L-Val-L-Tyr∼T_2_. The condensation domain (C) subsequently catalyzes peptide bond formation, which results in the dipeptide L-Val-L-Tyr∼S’T_2_, which is then reduced by the R-domain to an intermediate amino aldehyde that cyclizes to an imine. Oxidation to the pyrazinone final product likely occurs spontaneously [4].

Whereas in our analyses we used CDM containing 1 mM L-phenylalanine, 1 mM L-valine and 0.1 mM L-tyrosine, lower AA concentrations in the CDM were not tested. However, phevalin was shown to be produced by intracellular *S. aureus* [8] in experiments using the RPMI1640 medium (Thermo Fisher Scientific, Cat. No. 72400054), which contains 0.09 mM L-phenylalanine, 0.11 mM L-tyrosine and 0.17 mM L-valine, suggesting that reduced amino acid availability does not impact aureusimine production.

The BCAA valine is present in all three synthesized aureusimines and is the only natural substrate for the A_1_ domain [4]. BCAAs (isoleucine, leucine, valine) are essential metabolites and “expensive” goods for *S. aureus*. BCAA are both incorporated into membrane branched-chain fatty acids [18] where they comprise 65% of membrane fatty acids [19] and represent the most abundant amino acid in staphylococcal proteins [20].

In *S. aureus*, BCAA acquisition and biosynthesis are stringently monitored by the global regulator CodY [21], which uses isoleucine and GTP as co-factors [17, 22]. Transcription of genes involved in BCAA import (*brn*Q1, *brn*Q2, *bca*P), and biosynthesis (*leu* and *ilv* operons) is sequentially de-repressed upon drops in intracellular BCAAs [23, 24], with the bacterium prioritizing BCAA scavenging over *de novo* synthesis [17, 23]. *S. aureus* responds dramatically to valine depletion, as concentrations below 100 μM in the culture medium, lead to a prolonged lag phase (Supp. Fig 4D). Moreover, in the absence of exogenous valine, suppressor mutants in either *cod*Y or the 5’UTR of *ilv*D (first gene in the *ilv-leu* operon) are selected [17], highlighting the importance of valine availability for *S. aureus* growth. It is therefore remarkable that *S. aureus* uses valine as the main building block for secondary metabolites which are secreted at micromolar concentration.

While the NRPS A_1_ domain has a strict specificity for L-valine, A_2_ has a relaxed substrate specificity, for L-tyrosine, L-phenylalanine or, to a lesser extent, L-leucine. L-tyrosine, however, represents the preferred substrate [4].

The AAAs phenylalanine, tyrosine and tryptophan are synthesized from the central metabolite D-erythrose 4-phosphate, via the shikimate pathway [25]. Phenylalanine and tyrosine, similarly to BCAAs, are likely used for protein synthesis but not for catabolic purposes [28].The genes encoding the enzymes and proteins for the synthesis of AAAs (*trp, aro, tyr*) are controlled by repression via CodY, however, they are not as stringently repressed by CodY as genes involved in BCAA or histidine biosynthesis [23]. The delayed growth of *S. aureus* JE2 growth under deprivation of either AAAs or BCAAs (Supp. Fig 4), highlights the importance maintaining a stable intracellular pool for these metabolites.

AAAs are commonly known to be incorporated into secondary metabolites, however, a selective usage of exogenous vs. endogenous AAAs has not yet been observed. A possible explanation for exclusive usage of exogenous AAAs could be that *de novo* biosynthesis does not yield high enough amounts of phenylalanine or tyrosine, respectively, to ensure occupancy of the AusA A_2_ domains. The A_1_ domain, was shown to have a lower affinity to L-valine (K_M_= ∼3 mM), compared to the A_2_ domain affinity to either L-tyrosine (K_M_= ∼1) or L-phenylalanine (K_M_= ∼2). However, the specificity constants (K_cat_/K_M_ ratio) for A_1_ and A_2_ with their optimal substrates are nearly identical, indicating a balanced activity for both domains [4]. In wild-type *S. aureus de novo* biosynthesis yields ∼ 1.65 μmol/g pellet-associated valine upon growth in CDM [-Val], compared to ∼0.97 μmol/g upon growth in CDM.

Phenylalanine *de novo* biosynthesis seems to be less efficient, yielding only 0.05 μmol/g phenylalanine in CDM [-Phe], compared to 0.61 μmol/g during CDM cultivation. Therefore, reduced phenylalanine availability resulting from *de novo* biosynthesis could, in part, explain why phevalin remained undetectable under these conditions. Surprisingly, growth in CDM [-Tyr], yields comparable amounts of pellet-associated tyrosine as growth in CDM (0.03 vs 0.02 μmol/g, respectively), indicating that although this low concentration of tyrosine is sufficient for tyrvalin biosynthesis during growth in CDM, this is not the case for CDM [-Tyr]. This observation suggests that it is rather the source of tyrosine (exogenous vs endogenous) that impacts usage by AusA, than its concentration (Fig. 2I).

## CONCLUSION

Here, we utilize a metabolomics approach to show that in the AusA NRPS appears to fuel only exogenous phenylalanine and tyrosine into the cyclic dipeptides phevalin and tyrvalin, whereas the source of the BCAA valine can be either exogenous or endogenous.

Preferential usage of exogenous AAAs by AusA, can be explained by an association of the NRPS with the cytoplasmic membrane, in physical proximity to an AAA permease, to ensure an adequate supply of either of the two AAAs required for aureusimine biosynthesis. Other NRPSs, such as the pyoverdine NRPS enzymes in the Gram-negative *Pseudomonas aeruginosa* have been described to be associated with the cytoplasmic membrane and physically interact with precursor-generating enzymes to form membrane-bound multi-enzymatic complexes, as an efficient form of subcellular compartmentalization of a secondary metabolic pathway [26]. Additionally, a NRPS/polyketides synthase enzymatic complex involved in the biogenesis of the antibiotic bacillaene in the Gram-positive *B. subtilis* were shown to exhibit a similar membrane compartmentalization [27].

In summary, our observation suggests an unusual way of amino acid incorporation into secondary metabolites and opens new prospects for further explorations on NRPSs, both in *S. aureus* and other bacterial species.

## ACKNOWLEDGEMENTS

This work was supported by the Deutsche Forschungsgemeinschaft (DFG) in the project DFG RU 631/17-1 to T.R. and EI 384/16-1 to W.E. A. M. was partially supported by a grant of the German Excellence Initiative to the Graduate School of Life Sciences, University of Würzburg and the DAAD STIBET Fellowship of the University of Würzburg. We thank Wilma Ziebuhr and Christoph Schön (University of Würzburg) for providing various strains. We thank Peter Heinrich for initial experiments using chemically defined media. We thank Dr. Sebastian Blättner for cloning the pKOR1 *aus*AB knock-out plasmid.

## ETHICS DECLARATION

Not applicable

